# *Keetia nodulosa* sp. nov. (Rubiaceae - Vanguerieae) of West - Central Africa: bacterial leaf nodulation discovered in a fourth genus and tribe of Rubiaceae

**DOI:** 10.1101/2024.02.20.581267

**Authors:** Martin Cheek, Jean Michel Onana

## Abstract

*Keetia nodulosa* Cheek, a cloud forest climber nearly endemic to Cameroon, with a single record from Nigeria, is described and illustrated. It is remarkable as the first known species to be recorded with bacterial leaf nodules (BLN) in the genus *Keetia,* and also, in the tribe Vanguerieae. Other genera in Rubiaceae with BLN are *Psychotria* (Psychotrieae), *Sericanthe* (Coffeaeae) and *Pavetta* (Pavetteae). The BLN in *Keetia* (Vanguerieae) are illustrated for the first time here.

The characteristics and significance of bacterial leaf nodulation in *Keetia nodulosa* are discussed in the context of rapidly growing knowledge on the subject in flowering plants. *Keetia nodulosa* is provisionally assessed using the 2012 IUCN standard as Endangered (EN B2ab(iii). The importance of its conservation, and options for achieving this are discussed in the context of recent extinctions of other plant species in Cameroon.

This discovery of a new cloud forest species is discussed in relation to other cloud forest plant species described in the last twenty years which are also distributed over the highlands of the western half of Cameroon.

## Introduction

*Keetia* S.M. Phillips was segregated from *Canthium* Lam. by Bridson (1985, 1986). Restricted to sub-Saharan Africa, and extending from Guinea in West Africa (Gosline *et al*. 2023a; 2023b) to Sudan in the North and East (Darbyshire *et al*. 2015) and S. Africa in the South (Bridson 1986), this genus of about 40 accepted species (POWO, continuously updated) are mainly forest climbers, distinguished from similar Canthioid genera in west Africa by their pyrenes with a fully or partly-defined lid-like area around a central crest (Bridson 1986). In a phylogenetic analysis of the tribe based on morphology, nuclear ribosomal ITS and chloroplast *trnT-F* sequences, Lantz & Bremer (2004), found that based on a sample of four species, *Keetia* was monophyletic and sister to *Afrocanthium* (Bridson) Lantz & B. Bremer with strong support. Highest species diversity of *Keetia* is found in Cameroon and Tanzania, both of which have about 15 taxa (Onana 2011; POWO, continuously updated). In contrast, neighbouring Gabon has only 10 species, although most specimens recorded remain unidentified to species, Sosef *et al*. 2006). Several *Keetia* species are point endemics, or rare national endemics, and have been prioritized for conservation (e.g. Onana & Cheek 2011; Couch *et al*. 2019; Murphy *et al*. 2023; Darbyshire *et al*. 2023) and one threatened species, *Keetia susu* Cheek has a dedicated conservation action plan (Couch *et al*. 2022)

Bridson’s (1986) account of *Keetia* was preparatory to treatments of the Vanguerieae for the Flora of Tropical East Africa (Bridson & Verdcourt 1991) and Flora Zambesiaca (Bridson 1998). Pressed to deliver these, she stated that she could not dedicate sufficient time to a comprehensive revision of the species of *Keetia* outside these areas: “full revision of *Keetia* for the whole of Africa was not possible because the large number of taxa involved in West Africa, the Congo basin and Angola and the complex nature of some species would have caused an unacceptable delay in completion of some of the above Floras” (Bridson 1986). Further “A large number of new species remain to be described.” Several of these new species were indicated by Bridson (1986), and other new species by her arrangement of specimens in folders that she annotated in the Kew Herbarium. One of these species was later taken up and published by Jongkind (2002) as *Keetia bridsoniae* Jongkind. In the same paper, Jongkind discovered and published *Keetia obovata* Jongkind based on material not seen by Bridson. Based mainly on new material, additional new species of *Keetia* have been published by Bridson & Robbrecht (1993), Bridson (1994), Cheek (2006), Lachenaud *et al*. (2017), Cheek *et al*. (2018a) and Cheek & Bridson (2019).

In the course of formally publishing new species to science from Cameroon so that they could be Red Listed and considered for inclusion in the Cameroon Important Plant Areas programme (e.g. Murphy *et al*. 2023), numerous new species to science have been published (see below), mainly based on species informally identified as new in the course of a series of surveys for improved conservation management of plant species and habitats conducted mainly in western Cameroon in the 1990s (Cheek *et al*. 2006). This paper continues the endeavor.

In this paper, a remarkable new species of *Keetia, K. nodulosa* Cheek is described. *Keetia nodulosa* is unique in its genus and tribe for having conspicuous bacterial nodules on its abaxial leaf blade surfaces, resembling those seen in species of the genus *Pavetta* L., which also have conspicuous black nodules at nerve junctions. The presence of bacterial nodules was first reported in the conservation checklist “The Plants of Mount Kupe, Muanenguba and the Bakossi Mts” (Cheek *et al*. 2004: 375). The specimens *Etuge* 2798 and *Etuge* 2829 (both Mt Kupe) were matched with specimens from Cameroon, that had been included in the protologue of *Keetia purseglovei* Bridson (Bridson 1986), *Zenker* 2986 (Bipinde) and *Zenker & Staudt* 415 (Yaoundé). However, the two Etuge specimens concerned had been annotated as “vel sp. aff.”, indicating that they might represent another but related species. Further research showed that all the Ugandan material of *Keetia purseglovei,* including the type, lacked bacterial nodules, and while and filed similar to the Cameroonian material, differed in several morphological characters (see Table 1 below). In searching all other material of *Keetia* at K, and other herbaria, for bacterial nodules, an additional specimen, *Emwiogbon* FHI 65823 from Nigeria, close to the Cameroon border, was found. This matched the Cameroonian material of *K. nodulosa*. It had been identified as a second specimen of *Keetia inaequilatera* (Hutch. & Dalz.) Bridson. While similar to the type and only other known specimen of that species, characters were found that separated this specimen from the type of that species (see Table 1 and diagnosis below) including the presence (vs absence) of bacterial nodules. Finally, just before the paper was completed, a further specimen, with flower buds, *Gereau et al*. 5639 from the Rumpi Hills, that had been identified as *K. cf. hispida,* was encountered and also placed in *K. nodulosa* in view of having bacterial nodules and other concordant characteristics.

**Table 1.** Characters distinguishing *Keetia inaequilatera, K. nodulosa sp.nov*. and *Keetia purseglovei*. Data for the first and third species from Bridson (1986) and specimens at K.

|  | *Keetia inaequilatera* | *Keetia nodulosa* sp. nov. | *Keetia purseglovei* |
| --- | --- | --- | --- |
| Distribution | S.E. Nigeria | S.E. Nigeria & Cameroon | Uganda (and probably eastern DR Congo) |
| Habitat | Lowland forest <800 m alt. | Cloud forest 800 – 940 m alt. | Submontane forest 1200 – 1265 m alt. |
| Secondary (spur) shoots, number of nodes | (2 –)3 – 4(– 5) | 7 – 10(– 13) | 3 – 5 |
| Bacterial nodules on abaxial surface of leaf blades | Absent | Conspicuous along tertiary nerves | Absent |
| Domatia, position | On the secondary nerve bases | In the axils of the secondary nerves | In the axils of the secondary nerves |
| Domatia, number of hairs | 10 – 30 | 10 – 30 | 0 (–5) |
| Stipule persistence (fruiting stage) | Unknown | Highly caducous, present usually only at the distalmost node | Persistent for 3 – 4 nodes from the apex |
| Stipule blade shape at maturity | Triangular | Subquadrate | Transversely elliptic |
| Pedicel length (mm) | 2 – 3 | (1.8 –)2.5 – 3(–4) | 5 – 7 |
| Flower bud shape and surface | Capitate (apex ovoid, base narrow cylindric); papillate | Clavate (obovoid) to capitate; smooth | Constricted (waisted) in middle to capitate; smooth |
| Calyx indumentum | Glabrous | Apex of teeth densely long hairy (rarely glabrous) | Glabrous, or teeth with 2–3 hairs at apex |
| Leaf-blade shape (proximal leaves of spur shoots) and length: breadth ratio | Broadly ovate to suborbicular<br>1.2 – 1.5:1 | Narrow elliptic or obovate-elliptic<br>(2 –)3: 1 | Elliptic (rarely oblanceolate-elliptic)<br>2 – 2.5:1 |
| Leaf-blade colour, abaxial surface | Mid to dark brown | Grey-black (rarely green) | Pale orange or pale green |
| Number of secondary nerves on each side of the midrib | (3 –)4(– 5) | 5 – 7 | (4 –)5 – 6 |

Further searches on gbif.org revealed that additional specimens had been identified as *Keetia purseglovei*, mainly from Gabon, Central African Republic, R.D. Congo and Congo-Brazzaville. However, these differed from *K. nodulosa,* and only one of these, *Texier* 2164, possessed visible bacterial nodules (see notes below) so were discounted.

In this paper it is shown that two specimens from Cameroon previously ascribed to *Keetia purseglovei* in Bridson (1986) together with additional specimens, are specifically distinct from the Ugandan material of that species, including the type. The Cameroonian taxon, which extends to Nigeria, is formally characterized and named as *Keetia nodulosa* sp. nov.

Within Africa, Cameroon remains a major source of discovery for new species to science of vascular plants, with more species new to science published per annum than any other country in tropical Africa (Cheek *et al*. 2020a). Recent novelties range from forest trees (Quintanar *et al*. 2023; Cheek *et al*. 2022a; 2023a), shrubs and small trees (Couvreur *et al*. 2022; Gosline *et al*. 2022; Stone *et al*. 2023; Cheek *et al*. 2023b), lianas (Jongkind & Lachenaud 2022), rheophytes (Cheek *et al*. 2022b), terrestrial herbs (Cheek *et al*. 2021), to epilithic herbs (Janssens *et al*. 2022; Cheek *et al*. 2023c) and ferns (Shang & Zhang 2023; Dubuisson *et al*. 2022).

## Materials and Methods

Names of species and authors follow IPNI (continuously updated). Herbarium material was collected using the patrol method e.g. Cheek & Cable (1997). Identification and naming follows Cheek in Davies *et al*. (2023). Herbarium specimens were examined with a Leica Wild M8 dissecting binocular microscope fitted with an eyepiece graticule measuring in units of 0.025 mm at maximum magnification. The drawing was made with the same equipment with a Leica 308700 camera lucida attachment. Pyrenes were characterized by boiling selected ripe fruits for several minutes in water until the flesh softened and could be removed. Finally, a toothbrush was used to clean the exposed pyrene surface to expose the surface sculpture. Specimens were inspected from the following herbaria: BM, BR, K, P, WAG, YA. It was not possible to view the duplicates of *Keetia nodulosa* deposited at YA because they are thought to be in the mounting backlog (Onana pers. obs. Feb. 2024). The format of the description follows those in other papers describing new species of *Keetia*, e.g. Cheek & Bridson (2019). Terminology follows Beentje & Cheek (2013). Herbarium codes follow Index Herbariorum (Thiers, continuously updated). Nomenclature follows Turland *et al*. (2018). All specimens seen are indicated “!” The conservation assessment follows the IUCN (2012) standard.

## Results

### Taxonomic treatment

**Keetia nodulosa** *Cheek* **sp. nov.** Type: Cameroon, S.W. Province [now Region], Kupe-Muanenguba Division, alt. 850 m, Kupe Village, main trail towards Mount Kupe, forest near a valley, fr.16 July 1996, *Etuge* 2798 with Felix, Ewang, Bishop, P., Temple, R. (holotype K000109898!; isotypes BR0000025613452V!, MO, P, WAG1966136!, YA).

LSID: urn:lsid:ipni.org:names:77336635-1

*Keetia purseglovei* Bridson (1986: 972) *quoad Zenker* 2986 (BM!, BR!, K!, P!) and *Zenker & Staudt* 415 K!); Cheek *et al*. (2004: 375).

*Evergreen climber, climbing with clasping fruiting peduncles*, 5 – 10 m tall. Primary stems with distal internodes glabrous, drying purple at first, subglossy, longitudinally finely ridged (microscope needed), distal internode flattened, other internodes subterete with a small central hollow, 5.3 – 6.5 x 0.35 – 0.4 cm, (distal, fertile internodes) at length with epidermis becoming longitudinally streaked with white. *Secondary shoots* (brachyblasts, plagiotropic or spur shoots) leafy, opposite, subequal in pairs, each 12– 37 cm long, with 4 – 9 internodes, internodes 2.5 – 6.1 x 0.12 – 0.2.5(– 0.3) cm, otherwise as the primary stems (Fig. 1 **A**), glabrous at fruiting stage, at flowering stage with sparse, patent, bristle hairs as the leaves. *Leaves* of primary axis not seen; those of secondary shoots distichous, not dimorphic, opposite and equal at each node, thinly leathery to thickly papery, blades drying black on upper surface, grey-black, rarely grey-green, on lower surface, elliptic, narrowly elliptic, or obovate elliptic, (4.9 –)5.3– 8.5(– 10.8) x 2.3 – 3.9(– 4.7) cm, acumen triangular 0.4 – 1.1(– 1.5) x 0.25 – 0.5 cm long, apex rounded; base obtuse to broadly acute, or rounded, rarely subcordate, usually asymmetric and decurrent on petiole; midrib and secondary nerves dull white to pale yellow, raised on both surfaces; domatia pit-like, longitudinally elliptic-oblong, 0.4 – 0.55 x 0.25 mm, inserted in the axil of midrib and the subtending secondary nerve, with 14 – 25 copper-coloured bristle hairs c. 0.1 mm long inserted around the rim, directed randomly: inconspicuous on upper surface; margin slightly thickened, revolute; secondary nerves 5 – 6(– 7) on each side of the midrib, arising at 50– 60° from the midrib, curving gradually upwards, the apex terminating parallel to and 3 – 4 mm from the blade margin, sometimes uniting with the nerve above. Tertiary nerves faintly visible, quaternary nerves inconspicuous. *Bacterial nodules* conspicuous on the abaxial surface, jet black, mainly at the junctions or along the lengths of tertiary nerves, about (1 –)2 –4(–5) mm apart, each 0.75 –2.5 mm long, usually with 3 –7 short lateral lobes along their length, 0.3 mm (unlobed) or 0.5 –0.75 mm wide, with a few smaller, unlobed, T-shaped or comma shaped nodules interspersed (Fig. 1C); hairs sparse, 3 – 20 % cover, along the midrib, secondary nerves (abaxial surface), and margins, simple, pale bronze-coloured, 0.25 – 0.5 mm long, strigose, slightly curved from base to apex, distal part gradually tapering to an acute apex, leaf otherwise glabrous. *Petioles* plano-convex in transverse section, the adaxial surface extended as narrow wings, (0.4 –) 0.5 – 0.8(– 0.9) x 0.1 cm, indumentum as midrib of blade. *Stipules* free, caducous (usually persisting at terminal node only at fruiting stage), glabrous apart from colleters, at apical bud narrowly triangular, c. 8 x 2 mm, the blade not distinct from the awn; at older nodes (flowering stage only, Fig. 1D & E) the blade distinct, subquadrate, widest at base, 4.5 –5 x 4.5 –5 mm, midrib not conspicuous; awn excurrent from the outer surface of the bade, arising below the apex, c. 6 x 0.8 – 1 mm, terete (or longitudinally 5 ridged on both surfaces), apex acute; colleters inserted on the adaxial surface near the base, botuliform, 0.4 –0.8 x 0.2 mm, apex rounded. *Inflorescences* (Fig. 1A), axillary on spur (plagiotropic) branches, held above the stem, in 2 – 4 successive nodes beginning 1– (– 2) nodes below stem apex; anthesis ± simultaneously at all nodes, each inflorescence (11–) 40 – 60-flowered, forming heads 2.8 – 4.8 x 1.2 – 1.5 cm. Peduncles 15 – 22 x 0.75 mm, with two pairs of bracts 4 – 5 mm below the apex, bracts triangular c. 0.75 x 0.4 mm, membranous, sparsely and inconspicuously simple hairy; branches two, equal, 4 – 7 mm long, with each branch further forked, or terminating in a fascicle of 5 – 12 flowers. *Pedicels* 2.5 –3(–4) x 0.75 mm long, with several scattered, slightly spreading, straight, acute hairs 0.25 mm long. Calyx-hypanthium obconical 1 x 1.25 mm, with c. 5 shallow longitudinal grooves, calyx tube shortly cylindrical, 0.3 –0.4 mm long; teeth 5, very shortly and broadly triangular, 0.1– 0.2 x 0.5 mm, the margins of the teeth apices with dense erect, simple hairs 0.1 – 0.15 mm long as the pedicel, rarely absent, or, with a few on the abaxial surface (Fig. 1F&G). Corolla in bud clavate or narrowly obovoid, unconstricted 4.2 – 4.5(– 5) x 2 – 2.5 mm, apex rounded; at anthesis white, tube 3 x 2 mm, lobes 5, valvate, reflexed, oblong triangular, 2 x 1 –1.5 mm, mouth with exserted, moniliform white hairs 0.7 – 1.5 mm long, from a ring inserted 0.3 –0.4 mm below the mouth and 2 mm above the base of the corolla tube (Fig. 1F&K); inner surface glabrous from base to a ring of translucent deflexed bristle hairs c. 1.5 mm long adjacent to the ring of exserted hairs, inserted c. 2 mm above base (Fig. 1K). Stamens 5, inserted just below the corolla tube mouth, erect, filaments flat, 0.2 –0.3 x 0.2 mm (Fig. 1I); anthers exserted, introrse, narrowly ellipsoid, 1.5 x 0.5 mm, apical connective appendage conical, c. 0.1 – 0.1 mm (Fig. 1H&J), sub-basifixed, base minutely hastate, the two bases conical, splayed, c. 0.1 mm long, acute. Disc annular, truncate, c. 0.2 x 0.8 mm, puberulent, hairs c. 0.1 mm long (Fig. 1G). Style c. 9 mm long, 0.2 mm wide, terete, the apex with a narrowly cylindrical, 10-fluted head or *receptaculum pollinis*, c. 1.75 x 0.75 mm, stigmatic apex papillate. *Infructescences* 3 – 7(– 9)-fruited, peduncles, clasping, reflexing, axes glabrescent, with a few thinly scattered simple hairs 1 mm long (Fig. 1 **F.**). *Fruit* green (mature fruits), fleshy, didymous, in side view suborbicular, 10 (– 11) x 12 (– 13) x 11 – 13 x 8 mm, the two carpels united along their length but divided by a shallow longitudinal groove on each side (Fig. 1N), apex shallowly retuse, apical sinus c. 1 x 5 mm, including calyx 2 mm diam., teeth persistent, disc inconspicuous (Fig. 1 **G.**); base slightly cordate or rounded, surface with 2– 8 raised verrucae mainly on each side of each carpel, verrucae c. 1 x 1 mm; 1-seeded fruits (by abortion, the majority, 7/8 of all ripe fruit), ovoid-elliptic, asymmetric, (8 –)10– 11 x (7–)7.5– 8 mm. Pyrene pale brown, woody, subellipsoid, 0.9 – 1 x 0.7– 0.75 x 0.5 – 0.7 cm, the surface with low, irregular, orbicular raised areas c. 1 mm diam., interlaced with white fibres. Lid apical, cap-like, c. 2–3 x 6 x 6 mm, angled c. 20 degrees towards the ventral face, crest (keel) distinct, broad; ventral face sometimes with a transverse slit opening 2(–3) mm long, at junction with main body of pyrene. Fig. 1A-Q.

**Fig. 1.**
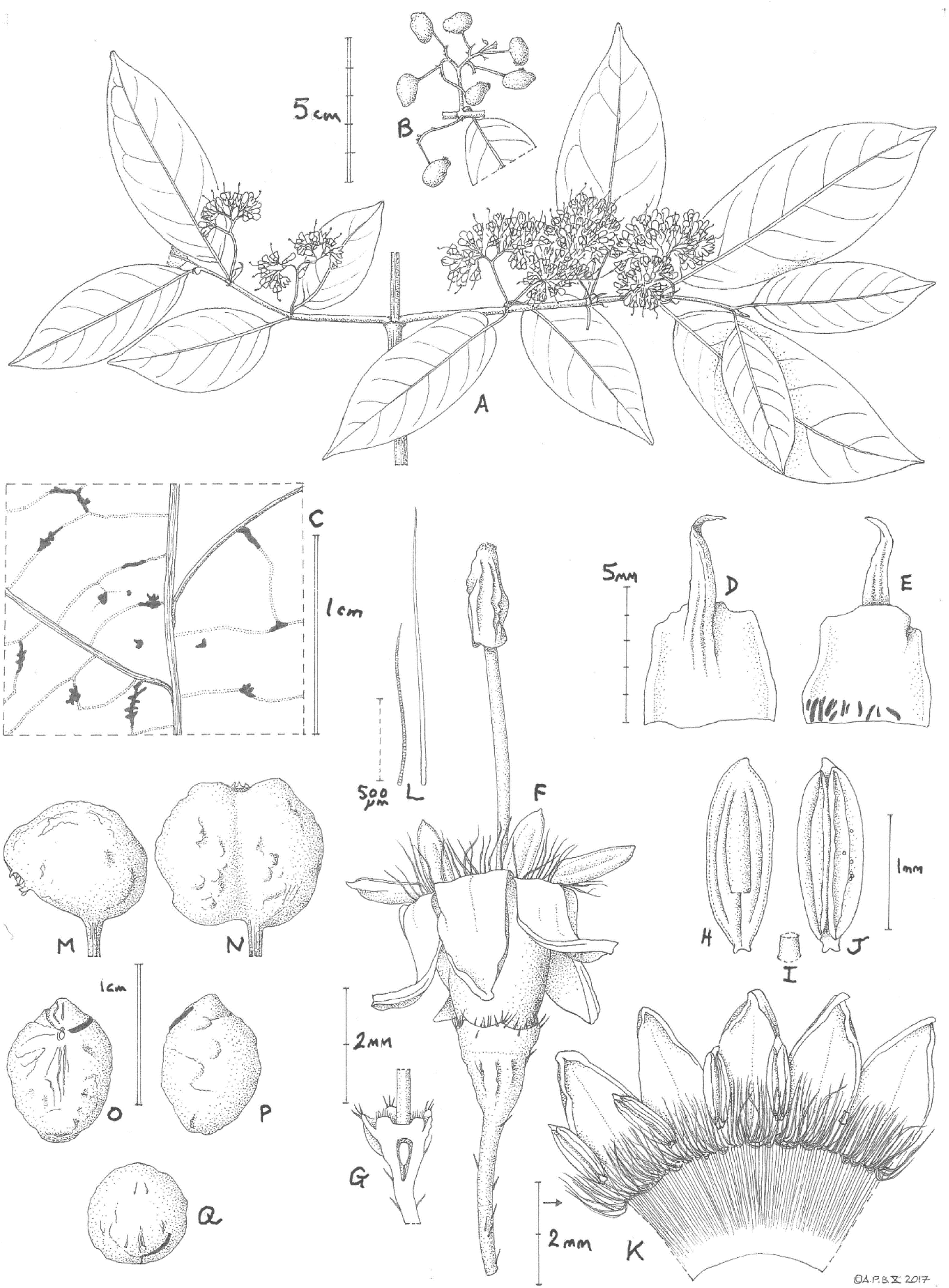
Keetia nodulosa. **A.** habit, flowering secondary (short plagiotropic or spur) shoots; **B.** infructescence; **C.** leaf-blade, abaxial surface showing bacterial nodules; **D.** stipule abaxial (outer) surface; **E.** stipule adaxial (inner) surface showing colleters; **F.** flower; **G.** near longitudinal section of flower base, showing disc; **H.** anther, outer surface; **I.** filament; **J.** anther, inner face; **K**. corolla, opened (one stamen removed); **L.** moniliform (L) and bristle (R) hairs from inner surface of corolla tube (see K); **M.** single seeded fruit; **N** double seeded fruit; **O** pyrene, frontal view; **P** pyrene, side view; **Q** pyrene, plan view. **A, C-L** from *Zenker* 415; **B, M-Q** from *Etuge* 2798. Drawn by ANDREW BROWN.

#### RECOGNITION

*Keetia nodulosa* differs from all known species of the genus in having bacterial nodules on the abaxial leaf blade surfaces (vs absent), further differing also from the similar *Keetia purseglovei* Bridson in the primary axis subterete (vs 4-fluted); stipules caducous at fruiting stage, persisting usually only at stem apex (vs persisting for 3 to 4 nodes from apex); stipule blades subquadrate (vs transversely elliptic); pedicels 2.5 –3(– 4) mm long (vs 5 – 7 mm) From *K. inaequilatera* (Hutch.) Bridson differing in the narrow elliptic or obovate-elliptic leaf blades with length: breadth ratio (2 –)3: 1(vs broadly ovate to suborbicular, 1.2 – 1.5:1), the domatia situated in the axils of the secondary nerves (vs on the secondary nerve bases) and the flower bud smooth, (not with the corolla bud head minutely papillate). See Table 1 above for additional diagnostic characters.

#### DISTRIBUTION

S.E. Nigeria and Cameroon

#### SPECIMENS EXAMINED

**NIGERIA.** South-Eastern State, Ikom District, Cross North Forest Reserve, Ikom. High forest, fr. 8 June 1972, *J.A. Emwiogbon* in FHI 65823 (FHI, K!). **CAMEROON. Central Region:** Yaoundé, “Yaunde Station**”** 800 m, fl. 1890–1894, *Zenker & Staudt* 415 (B destroyed; BM, K!); **South Region:** Bipinde, Urwaldgebiet, fr. 1904, *Zenker* 2986 (BM!, BR!, K!, P!); **South West Region**, Kupe Muanenguba Division, Kupe Village, main trail towards Mount Kupe, forest near a valley, fr. 16 July 1996, *Etuge* 2798 (holotype K(K000109898)!; iso. BR(BR0000025613452V)!, MO, P, WAG(1966136)!, YA); ibid, main trail towards Mount Kupe, 800 m alt., fr. 16 July 1996, *Etuge* 2829 (K!, YA); Ndian Division, Rumpi Hills, ca. 6 km E of Dikome Balue on foot path to Ifanga Nalende, ca., 300 m E of junction with trail to Momboriba, in primary forest on clay loam with *Garcinia* and *Coelocaryon* spp., buds 10 Dec.1994, *Gereau, F. Namata, E. Jato, E. Sarabe,* 5639 (K!, MO, YA)

#### HABITAT & ECOLOGY

Submontane evergreen forest (where known); 800 – 940 m alt. The altitudes of two of the specimens cited above (from Cross River North and from Bipinde) are not given on the label so it is possible that they are from lower altitudes than the other specimens, where altitude is recorded. However, both locations include points that exceed 800 m altitude, so it is conceivable that they are consistent with the remaining specimens in this respect.

#### PHENOLOGY

The initiation of flowering in December (dry season, *Gereau et al*. 5639) occurs at the same time as stem extension and is when new leaves are formed, while, when fruits are ripe (June and July, early wet season) the apical buds of the secondary stems appear dormant and there no new leaves are visible.

#### CONSERVATION STATUS

The relative frequency of occurrence of *Keetia nodulosa* is extremely low, indicating that even at its known locations it is extremely rare. At each location it is known from only a single collection, except at Mt Kupe where two collections are known. However, these two were collected on the same day, by the same team, on the same path up the mountain, and were 31 numbers apart. It may be that they were collected from the same plant, first in the morning on the way up, and then at the end of the day, on the way down. While at the Ikom location few collections of any plants have been made, at the Mt Kupe and Bipinde locations many thousands of herbarium specimens have been collected (e.g. Cheek *et al*. 2004), so if the species was not extremely rare, further records would be expected.

*Keetia nodulosa* is here provisionally assessed as Endangered (EN B2 ab(iii)) under the IUCN (2012) standard because five locations are known (see specimens examined above), each with observed or inferred imminent or actual threats of habitat clearance resulting from iron ore extraction infrastructure (Bipinde), quarrying and urbanisation (Yaoundé) and clearance for smallholder agriculture (Ikom, Rumpi Hills and Mt Kupe locations). *Keetia nodulosa* may already be extinct at the Yaoundé location due to the threats cited (Murphy *et al*. 2023). The area of occupation is assessed as 20 km^2^, using the IUCN required 4 km^2^ cell size. It is possible that the species also occurs in Gabon at Mt. Belinga (see notes below) but since the physical specimen, *Texier* 2164 has not been verified by the authors, only seen as an image (which shows some anomalous characters, see notes below), it is not included, taking the precautionary principle. *Keetia nodulosa* may yet be found in other locations within or outside the range documented here. However, the likelihood of this is not high, since tens of thousands of specimens have already been collected in surveys of suitable habitat in areas to the north and south of, and also within its known range (Cheek *et al*. 1992; Cheek *et al*. 1996; Cable & Cheek 1998; Cheek *et al*. 1996; 2000; Maisels *et al*. 2000; Chapman & Chapman 2001; Harvey *et al*. 2004; Cheek *et al*. 2004; Cheek *et al*. 2006; Cheek *et al*. 2010; Harvey *et al*. 2010; Cheek *et al*. 2011; Murphy *et al*. 2023).

#### ETYMOLOGY

The species is named for the bacterial nodules conspicuous on the abaxial leaf surfaces of this species, in which it is currently unique in the genus, and in the tribe.

#### PHENOLOGY

Flowering, new leaves and stem extension in December (dry season); fruiting June and July (early wet season).

### Notes

*Keetia nodulosa* is highly similar to *K. purseglovei*. The fruits, including the endocarps, which are so often useful in distinguishing species from each other in the genus, are more or less identical. It is not remarkable that material of the first species was included in the second.

In the protologue of *K. purseglovei*, five specimens from Cameroon are cited as paratypes, of which two are attributed here to *K. nodulosa* (see Specimens Examined above). A third Cameroonian paratype of *K. purseglovei, Bates* 1904 (Bitye, Ebolowa, BM! cited in error as 1940, but with a determination slip as *K. purseglovei* by Bridson) is a third, apparently undescribed species, differing from *K. nodulosa* in lacking bacterial nodules, in having suborbicular, strongly persistent stipules (vs subquadrate, caducous) and completely white, glossy primary stems (vs purple, streaked), and secondary stems completely glabrous in the flowering stage (vs sparse, patent, bristle hairs). This specimen also differs from *K. purseglovei s.s*. of Uganda, which has e.g. matt black primary stems, transversely elliptic mature stipules and much longer pedicels. *Bates* 1904 seems to represent yet another undescribed species. *Leeuwenberg* 5083 (60 km SW Eseka, BR image!, WAG image!) is a further paratype of *K. purseglovei*, also differing from *K. nodulosa* in lacking bacterial nodules. It appears to also differ from that species in lacking domatia, but this and other features needs to be confirmed by checking a physical specimen since even on the high quality images of BR, it is difficult to be certain. When this is possible, it may prove be conspecific with *Bates* 1904. Both specimens are lowland, c. 200 m alt. (vs 800 to 940 m alt. in *K. nodulosa*) and occur in southern Cameroon, between the Nyong and Ntem rivers. The remaining Cameroonian paratype, cited as *Bates* 1462 (BM) has not been found and neither has the remaining non Ugandan paratype of *K. purseglovei, Gossweiler* 9147 (BM, Zaire, Leopodville Province) (Cheek pers. obs. Jan. 2024).

Searching gbif.org for *Keetia purseglovei* retrieves 41 records of which 31 have associated images, and which amount to 18 unique specimen records. Apart from those attributable to *Keetia purseglovei* sensu stricto (Uganda, two specimens studied, also likely three specimens from DRC subject to confirmation after physical examination) and *K. nodulosa* (four specimens cited in this paper), specimens are also from Cameroon (*Leeuwenberg* and Bates see attributions above), the Central African Republic (3), Republic of Congo (1), Angola (1) and Gabon (2). Inspection of associated images, where available, and where resolution permits, reveals that with one exception, none have the bacterial nodules of *K. nodulosa*. These specimens also show dissimilarities with *Keetia purseglovei*. It is possible that they also may represent further new species to science, potentially conspecific with *Bates* 1904 (see above).

*Texier* 2164 (BR, BRLU, G, LBV, MO, P, WAG) was collected at the Mt Belinga chain, 60 km NE of Makoukou. The images available on gbif.org of plants live in the field clearly show black bacterial nodules on the abaxial surfaces of the leaves (https://www.tropi…imageid=100597044). Mt Belinga is known to host submontane forest, the habitat of *K. nodulosa*. However, *Texier* 2164 has 1) densely hairy stems, atypical of *K. nodulosa,* which has sparse hairs on the stem at the flowering stage, 2) leaf blades with length:breadth ratio c. 4: 1 (vs 2 – 3: 1), which 3) lack an acumen, 4) inflorescences 1.5 times the petiole length (vs c.3 – 4 times). Taken together these differences suggest that *Texier* 2164 may be a second species of *Keetia* with bacterial nodules. Verification of a physical specimen is desirable to establish a firm identification.

*Gereau et al*. 5639 had previously been identified (not by Bridson) as “*Keetia cf. hispida* sensu lato (aff. *setosum* Hiern)”. This was no doubt due to the setose hairs. However, *Keetia hispida s.l*. has swollen, ant-inhabited primary stem nodes, larger leaves with domatia along the secondary nerves, and lacks the quadrate stipule blades of *K. nodulosa. Gereau et al*. 5639 has only immature flower buds but is consistent with *K. nodulosa* in all features including the presence of bacterial nodules.

Variation within *Keetia nodulosa*. While the four fruiting specimens of *Keetia nodulosa* are relatively uniform morphologically, the sole specimen with open flowers, *Zenker & Staudt* 415 (“Yaunde-Station”) is slightly anomalous in that the leaves are longer (reaching 9 – 10 cm long vs <9 cm long). Only a small portion of one abaxial leaf is visible on the specimen, and this is insect-damaged, making unambiguous confirmation of the presence of bacterial nodules challenging. It is even possible that *Zenker & Staudt* 415 is taxonomically separable from the other specimens that comprise *Keetia nodulosa*

That bacterial nodules were not previously detected in specimens of *Keetia nodulosa*, two of which have been in herbaria for more than 100 years, is likely because there was no reason to expect them to be found. It was only the first author’s work identifying and describing other new species to science with bacterial nodules in the same location (Mt Kupe) and at about the same time (Cheek *et al*. 2008) that had raised awareness of this trait and facilitated its detection in the *Keetia* in 2004 (Cheek *et al*. 2004).

## Discussion

### Leaf bacterial nodulation

Since first reported (in *Pavetta,* Rubiaceae, Zimmerman 1902), knowledge of bacterial nodulation in leaves of flowering plants, occurring only in palaeotropical Primulaceae and Rubiaceae (but see notes on *Dioscorea* L. and *Stryrax* L. below), has been growing steadily. Reviews on the subject include Boodle (1923), Lersten and Horner (1976), Lemaire *et al*. (2011), Yang & Hu (2018), and Pinto-Carbó *et al*. (2018). ‘Bacterial leaf symbiosis’ is characterized as comprising endosymbiotic bacteria being organized in specialized leaf structures, usually known as nodules, or sometimes as galls, bacteriocecidia, or warts. These are visible macromorphological aspects of the symbiosis (Lemaire *et al*. 2011). The bacteria of the nodules are gram negative, rod or ellipsoid in shape, c. 2 micrometres long, and lack flagellae (Carlier *et al*. 2017). They are intercellular, and colonise the leaves through the stomata (Rubiaceae) or marginal teeth (Primulaceae) from the apical bud, from which inflorescences, flowers, and so eventually developing seeds, are also colonized.

The symbiotic bacteria concerned have been placed in the genus *Burkholderia* s.l. (Pinto-Carbó *et al*. (2018). Bacterial colonization of leaves without the bacteria being organized into visible leaf structures also occurs, with the bacteria thinly scattered inside the leaf (endophytic) between the mesophyll cells (Verstraete *et al*. 2017). Such endophytic non BLN bacteria occur more widely in genera of Rubiaceae than do BLN and have been reported from two non BLN genera of Coffeeae (Verstraete *et al*. 2023) and five non BLN genera of the Vanguerieae, but were not found in *Keetia* species sampled (Verstraete *et al*. 2013). Transmission of bacteria between plants is known to be mainly vertical (Pinto-Carbó *et al*. 2018). However, in the Rubiaceae, though not in Primulaceae, there is evidence that horizontal transfer can also occur (Pinto-Carbó *et al*. 2018). It is speculated that this is effected by sap sucking insects moving from plant to plant, since the guts of some of these insects are known to be home also to *Burkholderia* bacteria. Lemaire *et al*. (2011) is a detailed recent study on the taxonomic occurrence of bacterial leaf nodulation in host plants. It is focused on the phylogenetic placement (genes 16S, rDNA, *recA*, and *gyrB*) of the bacteria (endosymbionts) of 54 plant species in four of the six known leaf nodulated plant genera (see below). This amounts to nearly 10% of all known nodulated plant species. The genera *Ambylanthus* A.DC and *Ambylanthopsis* Mez, both Primulaceae of Asia in which BLN are recorded, were not sampled. The study confirmed that free living, soil dwelling bacteria are basal in the clade *Burkholderia* s.l. and sister to the leaf nodulating species. In almost all cases of BLN symbiosis, there is a 1:1 relationship of a species of bacteria with a taxon of plant. Only one example is known of a plant species, *Psychotria kirkii* Hiern, which has been colonised twice, by different taxa of bacteria (Lemaire *et al*. 2011). The earliest branching subclade of *Burkholderia* s.l. to colonise plants is that inhabiting some Asian *Ardisia* Sw. species (Primulaceae, formerly Myrsinaceae, Larson *et al*. 2023). The next earliest branching subclade colonises some species of the genus *Sericanthe* Robbr. (Rubiaceae Coffeeae, Cheek *et al*. 2018d), 11 to 12 of the 17 known species being considered to have nodules) and *Pavetta* (Rubiaceae Pavetteae De Block *et al*. 2015) of which 350/400 species are considered to have nodules). Another branch colonises several species of *Psychotria* L. (Rubiaceae, Psychotrieeae, Lachenaud 2019) in which c. 80/1400 species are nodulated. The penultimate branches colonise mainly further species of the genus *Pavetta* but include colonisation of some other species of both *Sericanthe* and *Psychotria*. The final subclades colonise the majority of the *Psychotria* BLN species. Thus, the genera *Psychotria, Pavetta,* and *Sericanthe* have each been colonized more than once, independently, by bacteria likely either from the soil or from other plants. Therefore, there have been multiple horizontal transfers of bacteria to leaf nodulated plant species, and co speciation or evolution of endosymbionts with their host plants through vertical transfer has not been universal. Divergence estimates by Lemaire *et al*. (2011) point to a relatively recent origin of bacterial symbiosis in Rubiaceae, dating back to the Miocene (5 to 23 Mya).

Following strong support from genome analysis, the bacterial genus *Burkholderia* s.l. has been divided into several genera which largely correspond to different lifestyles or symbioses (Estrada de los Santos *et al*. (2018). *Burkholderia* s.s. are human and animal pathogens, while symbionts of the fungal phytopathogen *Rhizopus microsporus* are now classified as *Mycetohabitans. Mimosa* root nodulating bacteria are classified as *Trinickia*, and ‘plant beneficial and environmental strains’ (including the bacterial nodulating leaf symbionts discussed above) are now classified as *Paraburkholderia*, which genus includes also other N_2_ fixing legume root symbionts. N_2_ fixing legumes are also colonized by bacteria of the genus *Caballeronia,* but *Caballeronia* are also endophytic in the leaves of the non BLN genera *Empogona* Hook.f. and *Tricalysia* A.Rich. ex DC. of Coffeaeae (Verstraete *et al*. 2023)*. Paraburkholderia* can also be symbionts of amoeba e.g. *Dictyostelium discoideum,* and of insect guts (Brock *et al*. 2020).

Bacterial leaf nodulation is also considered to occur in the tropical African monocot *Dioscorea sansibarensis* Pax (Dioscoreaceae), where folding of the leaf apices forms visible (pale green) pockets which allow development of bacterial colonies of *Orella dioscoreae* (Alcaligenaceae, Burkolderiales, Carlier *et al*. 2017). Bacterial colonisation of marginal leaf glandular hairs has been observed in *Styrax camporum* Pohl of Brazil (Styracaceae, Machado *et al*. 2014), but the bacteria, which are both intra and intercellular, remain unidentified and nodules are not formed.

The endosymbiont bacteria of Rubiaceae have a small genome size and low coding capacity, both characteristic of reductive genome evolution. Genome sizes range from 2.4 Mb to 6.1Mb, well below the c.8 Mb average of free living *Burkholderia* s.l. species. Loss of functional capacity likely explains the failure of repeated efforts to cultivate endosymbiont bacteria (Pinto-Carbó *et al*. 2018). Equally, cultivated plants which lack their endosymbionts grow poorly and eventually die (Verstraete *et al*. 2017).

Although the genome of endosymbionts is reduced, synthesis gene clusters have been detected in those of all *Psychotria* and *Pavetta* species investigated so far (Pinto-Carbó *et al*. 2018). Evidence that the novel C_7_N aminocyclitol kirkamide is synthesized by the symbiont bacteria in *Psychotria kirkii* is that while it is detected in leaves of plants with the endosymbiont, it is not in aposymbiotic plants (lacking the endosymbiont). The compound is toxic to arthropods and insects, suggesting a role in protecting the host against herbivory (Sieber *et al*. 2015). A related compound, streptol glucoside is also found in the nodulated leaves of the same species. It displays potent herbicidal activity and may have allelopathic properties (Pinto-Carbó *et al*. 2018). Prescence of such bacterial endosymbionts may thus be advantageous for the hosts and confer an evolutionary advantage over plants which lack such endosymbionts. We can hypothesise that because species with bacterial leaf nodules contain many more bacteria than non BLN species, the quantity of advantageous compounds produced by the bacteria might be higher, increasing the evolutionary advantage further.

The bacterial nodules in Rubiaceae vary in form from genus to genus, and also within genera. In *Psychotria* they usually black, raised and conspicuous to the naked eye on the abaxial leaf surface, scattered uniformly over the blade, the shape, size and density of the nodules helping to separate one species from another. In a minority of species the nodules are linear and positioned next to the midrib only (Lachenaud 2019; Cheek et al. 2008). In contrast, in *Pavetta,* the nodules are usually most conspicuous on the adaxial surface, also black but in other species green and inconspicuous unless viewed in transmitted light. Frequently they occur as thickenings at the junction of the tertiary nerves (Manning 1996). In *Sericanthe*, the nodules are often inconspicuous unless viewed in transmitted light, and often linear and arranged along the midrib (e.g. Sonké *et al*. 2012). The regular pattern and spacing of the nodules through the leaf identifies them as such and differentiates BLN from e.g. epidermal fungal colonies which are more localized to only part of a leaf.

In herbarium specimens of *Keetia nodulosa* the nodules have similarities with those commonly seen in *Pavetta* see above. They are black, conspicuous, slightly raised, and often at nerve junctions. However, they differ from most *Pavetta* in being conspicuous only abaxially, as in the BLN of *Psychotria*.

The discovery of bacterial nodules in a further tribe and genus of Rubiaceae was unexpected. A survey of the occurrence of endosymbiotic bacteria specifically in the Vanguerieae found that they only occur in five genera, in none of which are nodules formed, and none of which were *Keetia* (Verstraete *et al*. 2017).

Further work is needed to identify the species of bacterium that produces the nodules in *Keetia*. This can be done by genomic studies of dried leaf material (Danneels & Carlier 2023). The symbiont is almost certain to be a *Paraburkholderia,* given that all other leaf nodule forming endosymbionts of Rubiaceae belong to this genus, and that the non BLN endophytic bacteria recorded in Vanguerieae are also this genus (Verstraete *et al*. 2017). It will be especially interesting to find out in which subclade of *Paraburkholderia* it falls, and so to deduce the source and date of this colonization. It can be speculated that the colonization event is recent, since this is the only known nodule-forming species in a genus of 40 species. If the event was as old as in the other nodulated genera of Rubiaceae (see above), one might expect that a much higher number, and proportion of the species, would have been found to have been nodulated, as in those other three genera. We speculate that the event may have occurred in the vicinity of the Cross-Sanaga Interval (Cheek et al. 2001) which has the highest species and generic diversity per degree square in tropical Africa (Barthlott *et al*. 1996; Dagallier *et al*. 2020). Here, three of the five locations of *Keetia nodulosa* occur, two others being nearby). All three of the other Rubiaceae genera with bacterial nodules have centres of species diversity in the Cross-Sanaga Interval (Lachenaud 2019; Manning 1986; Sonke *et al*. 2012) from which horizontal transfer to *Keetia* mediated by sap-sucking insects may have occurred.

The discovery reported in this paper of bacterial leaf nodulation in a genus and tribe previously unknown to have this characteristic, is the first since the report 60 years ago by Petit (1962) of nodulation in some species he attributed to *Tricalysia* which are now placed in *Sericanthe*. It is conceivable that bacterial leaf nodulation remains to be found in other genera in which it is previously currently unknown.

### Submontane forest species in the western half of Cameroon

Additional rare, threatened species of submontane forest found with *Keetia nodulosa* at Mt Kupe, Rumpi Hills, or elsewhere within the range of the species are *Coffea montekupensis* Stoffel. (Rubiaceae, Stoffelen *et al*. 1997), *Psychotria hardyi* O.Lachenaud (Rubiaceae, Lachenaud 2019), *Memecylon kupeanum* R.D.Stone *et al*. (Melastomataceae, Stone *et al*. 2008), *Sabicea bullata* Zemagho *et al*. (Rubiaceae, Zemagho *et al*. 2014), *Impatiens frithii* Cheek (Balsaminaceae, Cheek & Csiba 2002), *Microcos magnifica* Cheek (Cheek 2017) and *Microcos rumpi* Cheek (Cheek *et al*. 2023a) both Malvaceae s.l./Grewiaceae, *Cola etugei* Cheek (Malvaceae s.l./Sterculiaceae, Cheek *et al*. 2020b), *Psychotria spp*. (Rubiaceae, Cheek *et al*. 2008), *Deinbollia oreophila* Cheek (Sapindaceae, Cheek & Etuge 2009), *Kupea martinetugei* Cheek (Triuridaceae, Cheek *et al*. 2003), and *Vepris zapfackii* Cheek (Rutaceae, Cheek & Onana 2021). In several cases the species were initially considered point endemics but were shown after further surveys, to be more widespread in the surviving cloud forests of western Cameroon. However, in other cases despite additional surveys, species have remained point endemics e.g. *Brachystephanus kupeensis* I.Darbysh. (Acanthaceae, Champluvier & Darbyshire 2009). The high level of endemism in these submontane forests (extending to Bioko) contributes to the high species and generic diversity levels reported in the Cross Sanaga Interval mentioned above. There is no doubt that additional species remain to be described from these forests, although further survey work is hampered by the secession struggle in the two anglophone Regions, South West and North West that began in December 2016 and has taken thousands of lives and displaced half a million people (https://en.wikipedia.org/wiki/Anglophone_Crisis, accessed Feb. 2024). South West Region contains the majority of the surviving submontane forest in western Cameroon, indeed in the whole of the Gulf of Guinea.

*Keetia nodulosa* is one of many new species to science that came to light partly or entirely as a result of surveys for conservation prioritization in Cameroon. Cameroon has the highest number of globally extinct plant species of all countries in continental tropical Africa (Humphreys *et al*. 2019). The extinction of species such as *Oxygyne triandra* Schltr. (Thismiaceae, Cheek *et al*. 2018b) and *Afrothisia pachyantha* Schltr. (Afrothismiaceae, Cheek & Williams 1999; Cheek *et al*. 2019; Cheek *et al*. 2023d) and at least two species of the African genus *Inversodicraea* (Cheek *et al*. 2017), are well known examples, recently joined by species such as *Vepris bali* Cheek (Rutaceae, Cheek *et al*. 2018c), *Vepris montisbambutensis* Onana (Onana & Chevillotte 2015) and *Ardisia schlechteri* Gilg (Murphy *et al*. 2023). However, another 127 potentially globally extinct Cameroon species have recently been documented (Murphy *et al*. 2023: 18 – 22).

It is critical now to detect, delimit and formally name species such as *Keetia nodulosa* as new to science, since until they are scientifically recognised, they are essentially invisible to science, and only when they have a scientific name can their inclusion on the IUCN Red List be facilitated (Cheek *et al*. 2020a). Most (77%) species named as new to science in 2023 are already threatened with extinction (Brown *et al*. 2023). Many new species to science have evaded detection until today because they are in genera that are long overdue full taxonomic revision as was the case with *Keetia nodulosa,* or because they have minute ranges which have remained unsurveyed until recently.

If further global extinction of plant species is to be avoided, effective conservation prioritization is crucial, backed up by investment in protection of habitat, ideally through reinforcement and support for local communities who often effectively own and manage the areas concerned. Important Plant Areas (IPAs) programmes, often known in the tropics as TIPAs (Darbyshire *et al*. 2017; Couch *et al*. 2019; Darbyshire *et al*. 2023; Murphy *et al*. 2023) offer the means to prioritize areas for conservation based on the inclusion of highly threatened plant species, among other criteria. Such measures are vital if further species extinctions are to be avoided of rare, highly threatened species such as *Keetia nodulosa*.

## Acknowledgements

Completion of work on this paper was through support of the Cameroon TIPAs programme from Players of People’s Postcode Lottery (PPL).

Ranee Prakash at BM (Natural History Museum, London) is thanked for facilitating access to that herbarium.

The authors thank the late Martin Etuge (see Murphy *et al*. 2023) for collecting the two specimens on Mt Kupe that initiated the writing of this paper. These specimens were collected with the support of volunteers and sponsored scientists arranged by Earthwatch Europe, Oxford.

Drs Satabié, Achoundong, Ngo Ngwe, Lagarde, and Tchiengue, the former and current directors, of IRAD (Institute of Research in Agronomic Development)-National Herbarium of Cameroon, and their staff are thanked for expediting the collaboration between our two institutes under the terms of our Memorandum of Collaboration.

Diane Bridson and an anonymous reviewer are thanked for constructive comments on an earlier draft of the paper. Shigeo Yasuda prepared and cleaned the endocarps ready for measuring.

The authors declare no conflict of interest.

## References

Barthlott, W., Lauer, W. & Placke, A. (1996). Global distribution of species diversity in vascular plants: towards a world map of phytodiversity. Erdkunde 50: 317 –327 10.1007/s004250050096

Beentje, H. & Cheek, M. (2003). Glossary. In: Beentje, H. (ed), Flora of Tropical East Africa. Balkema, Lisse.

Boodle, L. A. (1923). The Bacterial Nodules of the Rubiaceae. Bulletin of Miscellaneous Information (Royal Botanic Gardens, Kew), 1923(9), 346–348. 10.2307/4120240

Bridson, D. M. (1985). The reinstatement of *Psydrax* (Rubiaceae, subfam. Cinchonoideae tribe Vanguerieae) and a revision of the African species. Kew Bull. 40: 687 – 725. 10.2307/4109853

Bridson, D. M. (1986). The Reinstatement of the African genus *Keetia* (Rubiaceae, Cinchonoideae, Vanguerieae). Kew Bull. 41 (4): 956 – 994. 10.2307/4102996

Bridson D.M. (1994). A new species of *Keetia* (Rubiaceae-Vanguerieae) Kew Bull. 49: 803 – 807. 10.2307/4118075

Bridson D.M. (1998). Rubiaceae (Tribe Vanguerieae) Flora Zambesiaca 5(2): 1 – 377. 10.2307/4111186

Bridson, D. M. & Robbrecht, E. (1993). A spiny-fruited new *Keetia* (Rubiaceae, Vanguerieae) from Kivu (Zaire). Belg. Journ. Bot. 126: 29 – 32.

Bridson, D. M. & Verdcourt, B. (1991). Flora of Tropical East Africa – Rubiaceae, 3. Rotterdam/Brookfield, A.A.Balkema. 10.1201/9780203755860

Brock, D.A., Noh, S., Hubert, A.N., Haselkorn, T.S., DiSalvo, S., Suess, M.K., Bradley, A.S., Tavakoli-Nezhad, M., Geist, K.S., Queller, D.C. and Strassmann, J.E. (2020). Endosymbiotic adaptations in three new bacterial species associated with *Dictyostelium discoideum*: *Paraburkholderia agricolaris* sp. nov., Paraburkholderia hayleyella sp. nov., and Paraburkholderia bonniea sp. nov. PeerJ, 8, p.e9151. 10.7717/peerj.9151

Brown, M., Bachman, S., & Lughadha, E. N. (2023). Three in four undescribed plant species are threatened with extinction. New Phytologist. 10.1111/nph.19214

Cable, S. & Cheek, M. (1998). The Plants of Mt Cameroon, a Conservation Checklist. Royal Botanic Gardens, Kew.

Carlier, A., Cnockaert, M., Fehr, L., Vandamme, P., & Eberl, L. (2017). Draft genome and description of Orrella dioscoreae gen. nov. sp. nov., a new species of Alcaligenaceae isolated from leaf acumens of *Dioscorea sansibarensis*. Systematic and applied microbiology, 40(1), 11 – 21. 10.1016/j.syapm.2016.10.002

Champluvier, D. & Darbyshire, I. (2009). A revision of the genera *Brachystephanus* and *Oreacanthus* (Acanthaceae) in tropical Africa. Syst. & Geogr. Pl. 79: 115 – 192. 10.2307/25746.

Chapman, J. & Chapman, H. (2001). The Forests of Taraba and Adamawa States, Nigeria an Ecological Account and Plant Species Checklist. University of Canterbury: Christchurch, New Zealand. pp. 221.

Cheek, M. (2006). A New Species of *Keetia* (Rubiaceae-Vanguerieae) from Western Cameroon. Kew Bull. 61 (4): 591 – 594.

Cheek, M. (2017). *Microcos magnifica* (Sparrmanniaceae) a new species of cloudforest tree from Cameroon. PeerJ 5:e4137 10.7717/peerj.4137

Cheek M., Bridson D.M. (2019). Three new threatened *Keetia* species (Rubiaceae), from the forests of the Eastern Arc Mts, Tanzania. Gardens Bulletin Singapore 71(Suppl.2): 155 – 169. 10.26492/gbs71(suppl.2).2019-12

Cheek, M. & Cable, S. (1997). Plant Inventory for conservation management: the Kew-Earthwatch programme in Western Cameroon, 1993 – 96, pp. 29 – 38 in Doolan, S. (Ed.) African Rainforests and the Conservation of Biodiversity, Earthwatch Europe, Oxford.

Cheek, M. & Csiba, L. (2002). A new epiphytic species of *Impatiens* (Balsaminaceae) from western Cameroon. Kew Bull. 57: 669–674. 10.2307/4110997

Cheek, M. & Etuge, M. (2009). A new submontane species of *Deinbollia* (Sapindaceae) from Western Cameroon and adjoining Nigeria. Kew Bull. 64: 503–508. 10.1007/s12225-009-9132-4

Cheek, M. & Onana, J.M. (2021). The endemic plant species of Mt Kupe, Cameroon with a new Critically Endangered cloud-forest tree species, *Vepris zapfackii* (Rutaceae). Kew Bull 76: 721–734 10.1007/s12225-021-09984-x

Cheek, M. & Williams, S. (1999). A Review of African Saprophytic Flowering Plants. In: Timberlake, Kativu eds. African Plants. Biodiversity, Taxonomy & Uses. Proceedings of the 15th AETFAT Congress at Harare. Zimbabwe, 39 – 49.

Cheek, M., Achoundong, G., Onana, J-M., Pollard, B., Gosline, G., Moat, J., Harvey, Y.B. (2006). Conservation of the Plant Diversity of Western Cameroon. In: Ghazanfar SA, H.J. Beentje (eds). Proceedings of the 17th AETFAT Congress, Addis Ababa. Ethiopia, 779 – 791.

Cheek, M., Alvarez-Agiurre, M.G., Grall, A., Sonké, B., Howes, M-J.R., Larridon, L. (2018d). *Kupeantha* (Coffeeae, Rubiaceae), a new genus from Cameroon and Equatorial Guinea. PLoS ONE 13: 20199324. 10.1371/journal.pone.0199324

Cheek, M., S. Cable, F.N. Hepper, N. Ndam & J. Watts. (1996). Mapping plant biodiversity on Mt. Cameroon. pp. 110 – 120 in van der Maesen, van der Burgt & van Medenbach de Rooy (Eds), The Biodiversity of African Plants (Proceedings XIV AETFAT Congress). Kluwer. 10.1007/978-94-009-0285-5_16

Cheek, M., Corcoran, M. & Horwath, A. (2008). Four new submontane species of *Psychotria* (*Rubiaceae*) with bacterial nodules from Western Cameroon. Kew Bull 63, 405–418 10.1007/s12225-008-9056-4

Cheek, M., Darbyshire, I. & Onana, J.M. (2023b). Discovery and conservation of *Monanthotaxis bali* (Annonaceae) a new Critically Endangered (possibly extinct) montane forest treelet from Bali Ngemba, North West Region, Cameroon. Kew Bull 78, 259–270 10.1007/s12225-023-10117-9

Cheek, M., Edwards, S. & Onana, J.M. (2023a). A massive Critically Endangered cloud forest tree, *Microcos rumpi* (Grewiaceae) new to science from the Rumpi Hills, SW Region, Cameroon. Kew Bull 78, 247–258. 10.1007/s12225-023-10119-7

Cheek, M., Etuge, M. & Williams, S. (2019). *Afrothismia kupensis* sp. nov. (Thismiaceae), Critically Endangered, with observations on its pollination and notes on the endemics of Mt Kupe, Cameroon. Blumea 64: 158–164. 10.3767/blumea.2019.64.02.06

Cheek, M., Feika, A., Lebbie, A., Goyder, D., Tchiengue, B., Sene, O., Tchouto, P., van der Burgt, X. (2017). A synoptic revision of *Inversodicraea* (Podostemaceae). Blumea 62: 125 –156. 10.3767/blumea.2017.62.02.07

Cheek, M., Gosline, G. & Onana, J.M. (2018c). *Vepris bali* (Rutaceae), a new critically endangered (possibly extinct) cloud forest tree species from Bali Ngemba, Cameroon. Willdenowia 48: 285 – 292. 10.3372/wi.48.48207

Cheek, M., Harvey, Y.B., Onana, J-M. (2010). The Plants of Dom. Bamenda Highlands, Cameroon: A Conservation Checklist. Royal Botanic Gardens, Kew.

Cheek M, Harvey Y, Onana J-M. (2011). The Plants of Mefou Proposed National Park. Yaoundé, Cameroon: A Conservation Checklist. Royal Botanic Gardens, Kew.

Cheek, M., Hatt, S., & Onana, J. M. (2022a). *Vepris onanae* (Rutaceae), a new Critically Endangered cloud-forest tree species, and the endemic plant species of Bali Ngemba Forest Reserve, Bamenda Highlands Cameroon. Kew Bulletin, 1 – 15. 10.1007/s12225-022-10020-9

Cheek, M., Mackinder, B., Gosline, G., Onana, J.M., Achoundong, G. (2001). The phytogeography and flora of western Cameroon and the Cross River-Sanaga River interval. Systematics and Geography of Plants 71: 1097 – 1100. 10.2307/3668742

Cheek M., Magassouba, S., Molmou, D., Doré, T.S., Couch, C., Yasuda, S., Gore, C., Guest, A., Grall, A., Larridon, I., Bousquet, I.H., Ganatra, B., Gosline, G. (2018a). A key to the species of *Keetia* (Rubiaceae - Vanguerieae) in West Africa, with three new, threatened species from Guinea and Ivory Coast. Kew Bull. 73: 56. 10.1007/s12225-018-9783-0

Cheek, M., Molmou, D., Magassouba, S., Ghogue, J.-P. (2022b).Taxonomic Monograph of *Saxicolella* (Podostemaceae), African waterfall plants highly threatened by Hydro-Electric projects, with five new species. Kew Bull.,1 – 31. 10.1007/s12225-022-10019-2

Cheek, M., Nic Lughadha, E., Kirk, P., Lindon, H., Carretero, J., Looney, B., Douglas, B., Haelewaters, D., Gaya, E., Llewellyn, T., Ainsworth, M.,Gafforov, Y., Hyde, K., Crous, P., Hughes, M., Walker, B.E., Forzza, R.C., Wong, K.M., Niskanen, T. (2020a). New scientific discoveries: plants and fungi. *Plants*, People Planet 2: 371 – 388. 10.1002/ppp3.10148

Cheek, M., Onana, J-M., Pollard, B.J. (2000). The Plants of Mount Oku and the Ijim Ridge, Cameroon, a Conservation Checklist. Royal Botanic Gardens, Kew.

Cheek, M., Osborne, J., van der Burgt, X. et al. (2023c). *Impatiens banen* and *Impatiens etugei* (Balsaminaceae), new threatened species from lowland of the Cross-Sanaga Interval, Cameroon. Kew Bull 78, 67–82 10.1007/s12225-022-10073-w

Cheek, M., Pollard, B.J., Darbyshire, I, Onana, J.M. & Wild, C. (2004). The Plants of Kupe, Mwanenguba and the Bakossi Mts, Cameroon. A Conservation Checklist. Royal Botanic Gardens, Kew.

Cheek, M., Sidwell, K., Sunderland T. & Faruk, A. (1992). A Botanical Inventory of the Mabeta-Moliwe Forest. Royal Botanic Gardens, Kew; report to Govt. Cameroon from O.D.A.

Cheek, M., Soto Gomez, M., Graham, S. W., & Rudall, P. J. (2023d). Afrothismiaceae (Dioscoreales), a new fully mycoheterotrophic family endemic to tropical Africa. Kew Bulletin, 1 – 19. 10.1007/s12225-023-10124-w

Cheek, M., Tchiengue, B., Baldwin, I. (2020b). Notes on the plants of Bakossi, Cameroon, and the new Cola etugei and Cola kodminensis (Sterculiaceae-Malvaceae). Plant Ecology and Evolution 153: 108–119 10.5091/plecevo.2020.1662

Cheek, M., Tchiengué, B., van der Burgt, X. (2021). Taxonomic revision of the threatened African genus *Pseudohydrosme* Engl. (Araceae), with *P. ebo*, a new, critically endangered species from Ebo, Cameroon. PeerJ 9:e10689 10.7717/peerj.10689.

Cheek, M., Tsukaya, H., Rudall, P.J., Suetsugu, K. (2018b). Taxonomic monograph of Oxygyne (Thismiaceae), rare achlorophyllous mycoheterotrophs with strongly disjunct distribution. PeerJ 6: e4828. 10.7717/peerj.4828

Cheek M., Williams S. & Etuge M. (2003). *Kupea martinetugei*, a new genus and species of Triuridaceae from western Cameroon. Kew Bull. 58: 225–228. 10.2307/4119366

Couch, C., Cheek, M., Haba, P., Molmou, D., Williams, J., Magassouba, S., Doumbouya, S. and Diallo, M.Y., (2019). Threatened habitats and tropical important plant areas (TIPAs) of Guinea, West Africa. Kew: Royal Botanic Gardens, Kew. https://kew.iro.bl.uk/concern/books/ce6950c8-5ed7-4115-b6d4-c09a45b686ff?locale=en

Couch, C., Molmou, D., Magassouba, S., Doumbouya, S., Diawara, M., Diallo, M.Y., Keita, S.M., Koné, F., Diallo, M.C., Kourouma, S. and Diallo, M.B. (2022). Piloting development of species conservation action plans in Guinea. Oryx, pp.1 – 10. 10.1017/s0030605322000138

Couvreur, T.L.P., Dagallier, L.-P.M.J., Crozier, F., Ghogue, J.-P., Hoekstra, P.H., Kamdem, N.G., Johnson, D.M., Murray, N., Sonké, B. (2022). Flora of Cameroon 45 – Annonaceae. Phytokeys 207, 1 – 532. 10.3897/phytokeys.207.61432

Dagallier, L.P., Janssens, S.B., Dauby, G., Blach-Overgaard, A., Mackinder, B.A., Droissart, V., Svenning, J.C., Sosef, M.S., Stévart, T., Harris, D.J. & Sonké, B. (2020). Cradles and museums of generic plant diversity across tropical Africa. New Phytologist 225: 2196 – 2213. 10.1111/nph.16293

Danneels, B., Carlier, A. (2023). Whole-Genome Sequencing of Bacterial Endophytes From Fresh and Preserved Plant Specimens. In: Martin, F., Uroz, S. (eds) Microbial Environmental Genomics (MEG). Methods in Molecular Biology, vol 2605. Humana, New York, NY. 10.1007/978-1-0716-2871-3_7

Darbyshire, I. (continuously updated) Tropical Important Plant Areas. http://science.kew.org/strategic-output/tropical-important-plant-areas

Darbyshire, I., Anderson, S., Asatryan, A., Byfield, A., Cheek, M., Clubbe, C., Ghrabi, Z., Harris, T., Heatubun, C. D., Kalema, J., Magassouba, S., McCarthy, B., Milliken, W., Montmollin, B. de, Nic Lughadha, E., Onana, J.M., Saıdou, D., Sarbu, A., Shrestha, K. & Radford, E. A. (2017). Important Plant Areas: revised selection criteria for a global approach to plant conservation. Biodivers. Conserv. 26: 1767 – 1800. 10.1007/s10531-017-1336-6.

Darbyshire, I., Kordofani, M., Farag, I., Candiga, R. and Pickering, H. (2015). The Plants of Sudan and South Sudan. Royal Botanic Gardens, Kew.

Darbyshire, I., Richards, S., Osborne, J., Matimele, H., Langa, C., Datizua, C., Massingue, A., Rokni, S., Williams, J., Alves, T. and De Sousa, C., (2023). Important Plant Areas of Mozambique. Kew: Royal Botanic Gardens. https://kew.iro.bl.uk/concern/books/c60f1a8b-07b5-4a7a-9e7f-211b48586faf?locale=zh

Davies, N.M.J., Drinkell, C, Utteridge, T.M.A. (2023). The Herbarium Handbook. Kew Publishing

De Block, P., Razafimandimbison, S. G., Janssens, S., Ochoterena, H., Robbrecht, E., & Bremer, B. (2015). Molecular phylogenetics and generic assessment in the tribe Pavetteae (Rubiaceae). Taxon, 64(1), 79 – 95. 10.12705/641.19

Dubuisson, J. Y., Boucheron-Dubuisson, E., Le Péchon, T., Bausero, P., Droissart, V., Deblauwe, V., … & Rouhan, G. (2022). Diversity and taxonomy of the fern genus *Vandenboschia* Copel. (Hymenophyllaceae, Polypodiidae) in the Afro-Malagasy region and description of a new species. Botany Letters, 169(2), 268 – 283. 10.1080/23818107.2022.2062445

Estrada-de Los Santos, P., Palmer, M., Chávez-Ramírez, B., Beukes, C., Steenkamp, E.T., Briscoe, L., Khan, N., Maluk, M., Lafos, M., Humm, E. and Arrabit, M. (2018). Whole genome analyses suggests that *Burkholderia* sensu lato contains two additional novel genera (*Mycetohabitans* gen. nov., and *Trinickia* gen. nov.): implications for the evolution of diazotrophy and nodulation in the Burkholderiaceae. Genes, 9(8), p.389. 10.3390/genes9080389

Gosline, G., Bidault, E., van der Burgt, X. et al. (2023a). A Taxonomically-verified and Vouchered Checklist of the Vascular Plants of the Republic of Guinea. Sci. Data 10, 327 (2023). 10.1038/s41597-023-02236-6

Gosline, G. et al. (2023b). Checklist of the Vascular Plants of the Republic of Guinea– printable format (1.10). Zenodo. 10.5281/zenodo.7734985

Gosline, G., Cheek, M., Onana, J.M., Ngansop, Tchatchouang, E., van der Burgt, X.M., MacKinnon, L., Dagallier, L.M.J. (2022). *Uvariopsis dicaprio* (Annonaceae) a new tree species with notes on its pollination biology, and the Critically Endangered narrowly endemic plant species of the Ebo Forest, Cameroon. PeerJ. Jan 6;9:e12614. 10.7717/peerj.12614

Harvey, Y.B., Pollard, B.J., Darbyshire, I., Onana, J.-M., Cheek, M. (2004). The Plants of Bali Ngemba Forest Reserve. Cameroon: A Conservation Checklist. Royal Botanic Gardens, Kew.

Harvey, Y.B., Tchiengue, B., Cheek, M. (2010). The Plants of the Lebialem Highlands, a Conservation Checklist. Royal Botanic Gardens, Kew.

Humphreys, A.M., Govaerts, R., Ficinski, S.Z., Lughadha, E.N. and Vorontsova, M.S. (2019). Global dataset shows geography and life form predict modern plant extinction and rediscovery. Nature Ecology & Evolution 3.7: 1043 – 1047. 10.1038/s41559-019-0906-2

IPNI (continuously updated). The International Plant Names Index. http://ipni.org/ (accessed: 07/2023).

IUCN. (2012). *IUCN Red List Categories and Criteria*: Version 3.1. Second edition. – Gland, Switzerland and Cambridge, UK: IUCN. Available from: http://www.iucnredlist.org/ (accessed: April 2023).

Janssens, S.B., Taedoumg, H., Dessein, S. (2022) *Impatiens smetsiana*, another example of convergent evolution of flower morphology in *Impatiens*. Plant Ecology and Evolution 155(2): 248 – 260. 10.5091/plecevo.89701

Jongkind, C. H. & Lachenaud, O. (2022). Novelties in African Apocynaceae. Candollea, 77(1), 17 – 51. 10.15553/c2022v771a3

Jongkind, C. H. (2002). Two New Species of *Keetia* (Rubiaceae) from West Africa. Kew Bull. 57 (4): 989 – 992. 10.2307/4115730

Lachenaud, O. (2019). Revision of the genus *Psychotria* (Rubiaceae) in West and Central Africa: Volume 1. Opera Botanica Belgica 17.

Lachenaud, O. Luke, Q. Bytebier B. (2017). *Keetia namoyae* (Rubiaceae, Vanguerieae), a new species from eastern Democratic Republic of Congo. Candollea 72: 23 – 26. 10.15553/c2017v721a2

Lantz, H. & Bremer, B. (2004). Phylogeny inferred from morphology and DNA data: characterizing well-supported groups in Vanguerieae (Rubiaceae). Botanical Journal of the Linnean Society. 146: 257 – 283. 10.1111/j.1095-8339.2004.00338.x

Larson, D. A., Chanderbali, A. S., Maurin, O., Gonçalves, D. J., Dick, C. W., Soltis, D. E., … & Utteridge, T. M. (2023). The phylogeny and global biogeography of Primulaceae based on high-throughput DNA sequence data. Molecular Phylogenetics and Evolution, 182, 107702. 10.1016/j.ympev.2023.107702

Lemaire, B., Vandamme, P., Merckx, V., Smets, E., & Dessein, S. (2011). Bacterial leaf symbiosis in angiosperms: host specificity without co-speciation. PLoS One, 6(9), e24430. 10.1371/journal.pone.0024430

Lersten, N.R., Horner, H.T. (1976). Bacterial leaf nodule symbiosis in angiosperms with emphasis on Rubiaceae and Myrsinaceae. Bot. Rev 42, 145–214 10.1007/BF02860721

Machado, S. R., Teixeira, S. D. P., & Rodrigues, T. M. (2014). Bacterial leaf glands in Styrax camporum (Styracaceae): first report for the family. Botany, 92(5), 403 – 411. 10.1139/cjb-2013-0297

Maisels, F.M., Cheek, M., Wild, C. (2000). Rare plants on Mt Oku summit, Cameroon. Oryx 34: 136 – 140. 10.1017/s0030605300031057.

Manning, S. D. (1996). Revision of *Pavetta* subgenus *Baconia* (Rubiaceae: Ixoroideae) in Cameroon. Annals of the Missouri Botanical Garden, 87 – 150. 10.2307/2399970

Murphy, B., Onana, J.M., van der Burgt, X. M., Tchatchouang Ngansop, E., Williams, J., Tchiengué, B., Cheek, M. (2023). Important Plant Areas of Cameroon. Royal Botanic Gardens, Kew.

Onana, J.-M. (2011) The Vascular Plants of Cameroon. A Taxonomic Checklist with IUCN Assessments. Flore Du Cameroun 39. Ministry of Scientific Research and Innovation, Yaoundé, Cameroon.

Onana, J.M. & Cheek, M. (2011). Red Data Book of the Flowering Plants of Cameroon, IUCN Global Assessments. Royal Botanic Gardens, Kew.

Onana, J.M. & Chevillotte, H. (2015). Taxonomie des *Rutaceae – Toddalieae* du Cameroun revisitée: découverte de quatre espèces nouvelles, validation d’une combinaison nouvelle et véritable identité de deux autres espèces de *Vepris* Comm. ex A. Juss. Adansonia, sér. 3. 37: 103 – 129. c10.5252/a2015n1a7

Petit, E. (1962). Rubiaceae africanae IX: Notes sur les genres *Aidia*, *Atractogyne*, *Aulacocalyx*, *Batopedina*, *Gaertnera*, *Morinda*, *Mussaenda*, *Nauclea*, *Sabicea*, Schizocolea et Tricalysia. Bulletin du Jardin botanique de l’Etat à Bruxelles, 32(Fasc. 2), pp.173 – 198. 10.2307/3667228

Pinto-Carbó, M., Gademann, K., Eberl, L., & Carlier, A. (2018). Leaf nodule symbiosis: function and transmission of obligate bacterial endophytes. Current opinion in plant biology 44: 23 – 31. 10.1016/j.pbi.2018.01.001

POWO (continuously updated). Plants of the World Online. Facilitated by the Royal Botanic Gardens, Kew. http://www.plantsoftheworldonline.org/?f=accepted_names&q=Impatiens (downloaded 1 Dec. 2023).

Quintanar, A., Sonké, B., Simo-Droissart, M., Barberá, P., Libalah, M., Harris, D.J. (2023). A matter of warts: a taxonomic treatment for *Drypetes verrucosa* (Putranjivaceae, Malpighiales) and a new cauliflorous species from Cameroon and Nigeria, D. stevartii. Plant Ecology and Evolution 156(2): 160 – 173. 10.5091/plecevo.102004

Shang, H., & Zhang, L. B. (2023). Systematics of the Fern Genus *Didymochlaena* (Didymochlaenaceae). Systematic Botany, 48(1), 110 – 139. 10.1600/036364423X16758873924144

Sieber, S., Carlier, A., Neuburger, M., Grabenweger, G., Eberl, L., & Gademann, K. (2015). Isolation and total synthesis of kirkamide, an aminocyclitol from an obligate leaf nodule symbiont. Angewandte Chemie, 127(27), 8079 – 8081. 10.1002/ange.201502696

Sonke, B., Taedoumg, H., & Robbrecht, E. (2012). A reconsideration of the Lower Guinean species of *Sericanthe* (Rubiaceae, Coffeeae), with four new species from Cameroon and Gabon. Botanical Journal of the Linnean Society, 169(3), 530 – 554. 10.1111/j.1095-8339.2012.01254.x

Sosef, M.S.M., Wieringa, J.J., Jongkind, C.C.H., Achoundong, G., Azizet Issembé, Y., Bedigian, D., Van Den Berg, R.G., Breteler, F.J., Cheek, M., Degreef, J. (2006). Check-list des plantes vasculaires du Gabon. Scripta Botanica Belgica 35. National Botanic Garden of Belgium. 435 pp.

Stoffelen, P., Cheek, M., Bridson, D., Robbrecht, E. (1997). A new species of *Coffea* (Rubiaceae) and notes on Mt Kupe (Cameroon). Kew Bulletin 52: 989–994. 10.2307/4117826

Stone, R. D., Ghogue J.-P. & Cheek, M. (2008). Revised treatment of *Memecylon* sect. *Afzeliana* (*Melastomataceae-Olisbeoideae*), including three new species from Cameroon. Kew Bull. 63: 227–241. 10.1007/s12225-008-9033-y

Stone, R.D., Tchiengué, B., Cheek, M. (2023). The endemic plant species of Ebo Forest, Littoral Region, Cameroon with a new Critically Endangered cloud forest shrub, Memecylon ebo (Melastomataceae-Olisbeoideae). bioRxiv 12.20.572583 10.1101/2023.12.20.572583

Thiers, B. (continuously updated). Index Herbariorum: A global directory of public herbaria and associated staff. New York Botanical Garden’s Virtual Herbarium. [continuously updated]. – Available from: http://sweetgum.nybg.org/ih/ (accessed: Feb. 2023).

Turland, N.J., Wiersema, J.H., Barrie, F. R., Greuter, W., Hawksworth, D.L., Herendeen, P.S., Knapp, S., Kusber, W.-H., Li, D.-Z., Marhold, K., May, T.W., McNeill, J., Monro, A.M., Prado, J., Price, M.J. & Smith, G.F. (ed.) (2018). International Code of Nomenclature for algae, fungi, and plants (Shenzhen Code) adopted by the Nineteenth International Botanical Congress Shenzhen, China, July 2017. – Glashütten: Koeltz Botanical Books. [= Regnum Veg. **159**]

Verstraete, B., Janssens, S., Lemaire, B., Smets, E., & Dessein, S. (2013). Phylogenetic lineages in Vanguerieae (Rubiaceae) associated with *Burkholderia* bacteria in sub-Saharan Africa. American Journal of Botany 100(12), 2380–2387. 10.3732/ajb.1300303

Verstraete, B., Janssens, S., & Rønsted, N. (2017). Non-nodulated bacterial leaf symbiosis promotes the evolutionary success of its host plants in the coffee family (Rubiaceae). Molecular Phylogenetics and Evolution, 113, 161 – 168. 10.1016/j.ympev.2017.05.022

Verstraete, B., Janssens, S., De Block. P., Asselman, P., Méndez, G., Ly, S., Hamon, P., Guyot, R. (2023). Metagenomics of African *Empogona* and *Tricalysia* (Rubiaceae) reveals the presence of leaf endophytes. PeerJ 11:e15778 10.7717/peerj.15778

Yang, C. J., & Hu, J. M. (2018). Bacterial leaf nodule symbiosis in flowering plants. Symbiosis, 2, 1 – 24. 10.5772/intechopen.73078

Zemagho, L., Lachenaud, O., Dessein, S., Liede-Schumann, S. & Sonké, B. (2014). Two new *Sabicea* (Rubiaceae) species from West Central Africa: *Sabicea bullata* and *Sabicea urniformis*. Phytotaxa, 173(4), 285 – 292. 10.11646/phytotaxa.173.4.3

Zimmermann, A. (1902). Über Bakterienknoten in den Blättern einiger Rubiaceen. Jahrbücher für wissenschaftliche Botanik 37: 1 – 11.

